# Perinatal THC Exposure via Lactation Induces Lasting Alterations to Social Behavior and Prefrontal Cortex Function in Rats at Adulthood

**DOI:** 10.1101/2020.02.03.931733

**Authors:** Andrew F Scheyer, Milene Borsoi, Olivier JJ Manzoni

**Affiliations:** INMED, INSERM U1249, Marseille, France; Aix-Marseille University, France; Cannalab Cannabinoids Neuroscience Research International Associated Laboratory. INSERM-Aix-Marseille University/Indiana University

## Abstract

Cannabis is the world’s most widely abused illicit drug and consumption amongst women during and surrounding the period of pregnancy is increasing. Previously, we have shown that cannabinoid exposure via lactation during the early postnatal period disrupts early developmental trajectories of prefrontal cortex maturation and induces behavioral abnormalities during the first weeks of life in male and female rat progeny. Here, we investigated the lasting consequences of this postnatal cannabinoid exposure on synaptic and behavioral parameters in the adult offspring of Δ9-tetrahydrocannabinol (THC)-treated dams. At adulthood, these perinatally THC-exposed rats exhibits deficits in social discrimination accompanied by an overall augmentation of social exploratory behavior. These behavioral alterations were further correlated with multiple abnormalities in synaptic plasticity in the prefrontal cortex, including lost endocannabinoid-mediated long-term depression (LTD), lost long-term potentiation and augmented mGlu2/3-LTD. Finally, basic parameters of intrinsic excitability at prefrontal cortex pyramidal neurons were similarly altered by the perinatal THC exposure. Thus, perinatal THC exposure via lactation induces lasting deficits in behavior and synaptic function which persist into adulthood life in male and female progeny.

## Introduction

Cannabis is currently the most widely used illicit drug in the world, and its consumption by pregnant women has been steadily increasing ^1–4^. Further, the majority of pregnant women estimate little-or-no risk associated with cannabis use during pregnancy ^5^, and a trend towards recommendations for cannabis use during the perinatal period has been noted in areas of the United States where cannabis use has been legalized ^6^. During the early postnatal period, breastfeeding by cannabis-using mothers actively transfers the principle psychoactive component of cannabis, Δ9-tetrahydrocannabinol (THC) and other cannabinoids to the developing infant, as the lipophilic nature of these cannabinoids results in concentrations in breast milk which are equal to or exceeding that found in the plasma of the user ^7,8^. Importantly, this transfer may persist well beyond the intoxication period of the user due to the long half-life of THC in breast milk ^9^, thus occluding the possibility of simply breastfeeding outside of the acute cannabis consumption period as is often done with alcohol consumption ^10^. Therefore, understanding and disseminating data regarding the consequences of THC consumption during the breastfeeding period is paramount to public health in the evolving age of cannabis use.

During prenatal and early postnatal development, herein referred to as the perinatal period (further defined in ^11,12^), the developing brain is acutely sensitive to exogenous cannabinoids. Indeed, the endocannabinoid system (ECS), comprised of the cannabinoid receptors (CB1R and CB2R), as well as the endogenous cannabinoids anandamide (AEA) and 2-arachidonoylglycerol (2-AG) amongst other elements, is intimately involved in development beginning at ontogenesis ^13,14^ and continuing throughout the neurodevelopmental period ^15–19^. Thus, it is not surprising that in utero exposure to THC induces wide-ranging, multi-scale lasting effects in a variety of brain regions and behavioral domains ^20^.

One such impacted brain region is the prefrontal cortex (PFC), a functional hub which plays a central role in such diverse cognitive realms as working memory, reasoning, cognitive flexibility and emotionally-guided behaviors including social interactions ^21,22^. The PFC is also a site of dense expression of CB1R ^23^ and is exquisitely sensitive to various ECS-related synaptopathies which manifest as a variety of cognitive disease states ^24,25^. Previously, our lab has demonstrated that exogenous cannabinoid exposure during in utero development results in significant abnormalities in PFC synaptic function which underlie behavioral alterations lasting into adulthood in cannabinoid-exposed progeny ^26,27^.

During the early postnatal period, we have shown that exposure to exogenous cannabinoids via lactation also alters the developmental trajectory of the PFC ^28^. Specifically, the timeline of cell excitability changes mediated by GABAergic maturation is delayed by exposure to either THC or a synthetic agonist of CB1R. This delayed PFC development is accompanied by select behavioral abnormalities which similarly indicate a retarded maturational profile in the cannabinoid-exposed offspring. Here, we extend upon these findings by investigating the long-term consequences of this perinatal THC exposure. We observed that, at adulthood (>90 postnatal days; PND), the offspring of dams treated with THC from PND1-10 exhibit augmented social behavior at the cost of social memory discrimination, as well as altered multi-directional synaptic plasticity and cell-excitability in the PFC. These data illuminate significant consequences resultant of perinatal THC exposure via lactation which may serve to inform breastfeeding mothers during the early postnatal period and provide fundamental bases for strategic interventions to correct such effects in impacted progeny.

## Materials and Methods

Further information and requests for resources/reagents should be directed to the Lead Contact, Olivier J.J. Manzoni.

### Animals

Animals were treated in compliance with the European Communities Council Directive (86/609/EEC) and the United States NIH Guide for the Care and Use of Laboratory Animals. The French Ethical committee authorized th’ project “Exposition Périnatale aux cannabimimétiques” (APAFIS#18476-2019022510121076 v3). All rats were group-housed with 12h light/dark cycles with ad libitum access to food and water. All behavioral, biochemical and synaptic plasticity experiments were performed on male and female RjHan:wi-Wistar rats (>P90) from pregnant females obtained from Janvier Labs. Pregnant dams arrived at E15 and remained undisturbed until delivery. Newborn litters found before 05:00p.m. were considered to be born that day (P0).

Dams were injected daily sub-cutaneously (s.c.) from P01-10 with Δ9-Tetrahydrocannabinol (THC; 2mg/kg/day). THC was suspended in 1:1:18 DMSO, cremophor and saline, and injected at 1ml/kg. Control dams (Sham) received vehicle.

### Behavior

Open field observations were conducted after rats were adapted to the room laboratory conditions for at least 1 h prior to testing. Tests were conducted in a 45 × 45 cm Plexiglass arena with ±2 cm of wood shavings covering the floor. All behavioral procedures were performed between 10:00 am and 3:00 pm. All sessions were recorded using a video camera using the Ethovision XT 13.0 video tracking software (Noldus, Netherlands) and analyzed by a trained observer who was unaware of treatment condition.

The social interaction apparatus consisted of a transparent acrylic chamber (120 × 80 cm) divided into three equal compartments (40 cm each) partially separated by white walls. The central compartment was empty and lateral compartments had an empty wire cage (20 cm diameter) were an object or a new rat (social stimulus) were placed during the test.

THC- or sham-exposed rats were individually habituated to the test cage containing the two empty wire cages for 10 min immediately prior to testing. The first trial (social approach, 5 min duration) consisted of giving the tested rat the option to socialize with either a novel object or a new, naïve, age- and sex-mate conspecific rat that were placed into the wire cages positioned on the arena’s opposite sides. Thirty minutes later, the tested rat returned to the apparatus for the second trial (social memory, 5min duration) wherein the two compartments held either the now-familiar rat from the first testing phase or a second, previously unknown, naïve, age- and sex-mate conspecific.

Only rats with no compartment preference during the habituation phase were used.

Time spent in each compartment and time spent exploring wire cages during the social approach and social memory phases were scored. Social Preference Ratio was calculated as time spent exploring either the wire cage containing the object, or the new rat divided by total time exploring both wire cages. Likewise, Social Memory Ratio was calculated as time spent exploring either the wire cage containing the rat used in the first trial or the new rat divided by total time exploring both wire cages.

### Slice preparation

Adult male and female rats were anesthetized with isoflurane and killed as previously described ^26,27,29,30^. The brain was sliced (300 mm) in the coronal plane with a vibratome (Integraslice, Campden Instruments) in a sucrose-based solution at 4°C (in mm as follows: 87 NaCl, 75 sucrose, 25 glucose, 2.5 KCl, 4 MgCl_2_, 0.5 CaCl_2_, 23 NaHCO_3_ and 1.25 NaH_2_PO_4_). Immediately after cutting, slices containing the medial prefrontal cortex (PFC) were stored for 1 hr at 32°C in a low-calcium ACSF that contained (in mm) as follows: 130 NaCl, 11 glucose, 2.5 KCl, 2.4 MgCl_2_, 1.2 CaCl_2_, 23 NaHCO_3_, 1.2 NaH_2_PO_4_, and were equilibrated with 95% O_2_/5% CO_2_ and then at room temperature until the time of recording. During the recording, slices were placed in the recording chamber and superfused at 2 ml/min with low Ca^2+^ ACSF. All experiments were done at 32°C. The superfusion medium contained picrotoxin (100 mM) to block gamma-aminobutyric acid types A (GABA-A) receptors. All drugs were added at the final concentration to the superfusion medium.

### Electrophysiology

Whole cell patch-clamp of visualized layer five pyramidal neurons mPFC and field potential recordings were made in coronal slices containing the mPFC as previously described ^31^. Neurons were visualized using an upright microscope with infrared illumination. The intracellular solution was based on K+ gluconate (in mM: 145 K+ gluconate, 3 NaCl, 1 MgCl_2_, 1 EGTA, 0.3 CaCl_2_, 2 Na2+ ATP, and 0.3 Na^+^ GTP, 0.2 cAMP, buffered with 10 HEPES). To quantify the AMPA/NMDA ratio we used a CH_3_O_3_SCs-based solution (in mM: 128 CH_3_O_3_SCs, 20 NaCl, 1 MgCl_2_, 1 EGTA, 0.3 CaCl_2_, 2 Na^2+^ ATP, and 0.3 Na^+^ GTP, 0.2 cAMP, buffered with 10 HEPES, pH 7.2, osmolarity 290 – 300 mOsm). The pH was adjusted to 7.2 and osmolarity to 290 – 300 mOsm. Electrode resistance was 4 – 6 MOhms.

Whole cell patch-clamp recordings were performed with an Axopatch-200B amplifier as previously described ^26,31–33^. Data were low pass filtered at 2kHz, digitized (10 kHz, DigiData 1440A, Axon Instrument), collected using Clampex 10.2 and analyzed using Clampfit 10.2 (all from Molecular Device, Sunnyvale, USA).

A −2 mV hyperpolarizing pulse was applied before each evoked EPSC in order to evaluate the access resistance and those experiments in which this parameter changed >25% were rejected. Access resistance compensation was not used, and acceptable access resistance was <30 MOhms. The potential reference of the amplifier was adjusted to zero prior to breaking into the cell. Cells were held at −75mV.

Current-voltage (I-V) curves were made by a series of hyperpolarizing to depolarizing current steps immediately after breaking into the cell. Membrane resistance was estimated from the I– V curve around resting membrane potential ^29^.

Field potential recordings were made in coronal slices containing the mPFC or the accumbens as previously described ^31^. During the recording, slices were placed in the recording chamber and superfused at 2 ml/min with low Ca^2+^ ACSF. All experiments were done at 32°C. The superfusion medium contained picrotoxin (100 mM) to block GABA Type A (GABA-A) receptors. All drugs were added at the final concentration to the superfusion medium. The glutamatergic nature of the field EPSP (fEPSP) was systematically confirmed at the end of the experiments using the ionotropic glutamate receptor antagonist CNQX (20 mM), which specifically blocked the synaptic component without altering the non-synaptic.

Both fEPSP area and amplitude were analyzed. Stimulation was performed with a glass electrode filled with ACSF and the stimulus intensity was adjusted ∼60% of maximal intensity after performing an input–output curve (baseline EPSC amplitudes ranged between 50 and 150 pA). Stimulation frequency was set at 0.1 Hz.

### Data acquisition and analysis

The magnitude of plasticity was calculated at 0-10min and 35–40 min after induction (for TBS-LTP and eCB-LTD) or drug application (mGlu2/3-LTD) as percentage of baseline responses. Statistical analysis of data was performed with Prism (GraphPad Software) using tests indicated in the main text after outlier subtraction (Grubb’s test, alpha level 0.05). All values are given as mean ±SEM, and statistical significance was set at p<0.05.

## Results

### Perinatal THC exposure augments social exploration at the cost of discrimination between novel and familiar social stimuli

In rodent models, exposure to cannabinoids (both synthetic and plant-derived) during gestation or early development induces an array of deleterious consequences on behavior manifesting both at early life and adulthood ^20,26–28,34,35^. Previously, we have shown that the administration of cannabinoids to lactating dams during the early postnatal period causes significant delays in both synaptic and behavioral development ^28^. Here, we used the same protocol of perinatal cannabinoid exposure in order to determine if such early-life exposure results in consequences lasting into adulthood. Thus, lactating rat dams were treated with a single daily low dose of Δ9-Tetrahydrocannabinol (THC; 2mg/kg, S.C.) or its vehicle (herein referred to as “sham”) from postnatal day (PND) 1-10. Experiments were conducted in both sexes of their adult offspring (>PND90).

First, in order to determine if simple parameters of naturalistic behavior were altered in the adult offspring of dams exposed to THC during the early postnatal period, we tested these animals in the open field environment (Figure 1a-c). Here, we did not observe any significant differences between the THC-exposed animals and those originating from sham-treated dams. Specifically, no differences were found in the time spent in the center of the arena, the total distance covered during the test, or the number of rearing events. Thus, perinatal THC exposure via lactation does not appear to alter baseline levels of spontaneous activity or anxiety at adulthood. Sex-specific details can be found in Tables 1-5.

**Figure 1.**
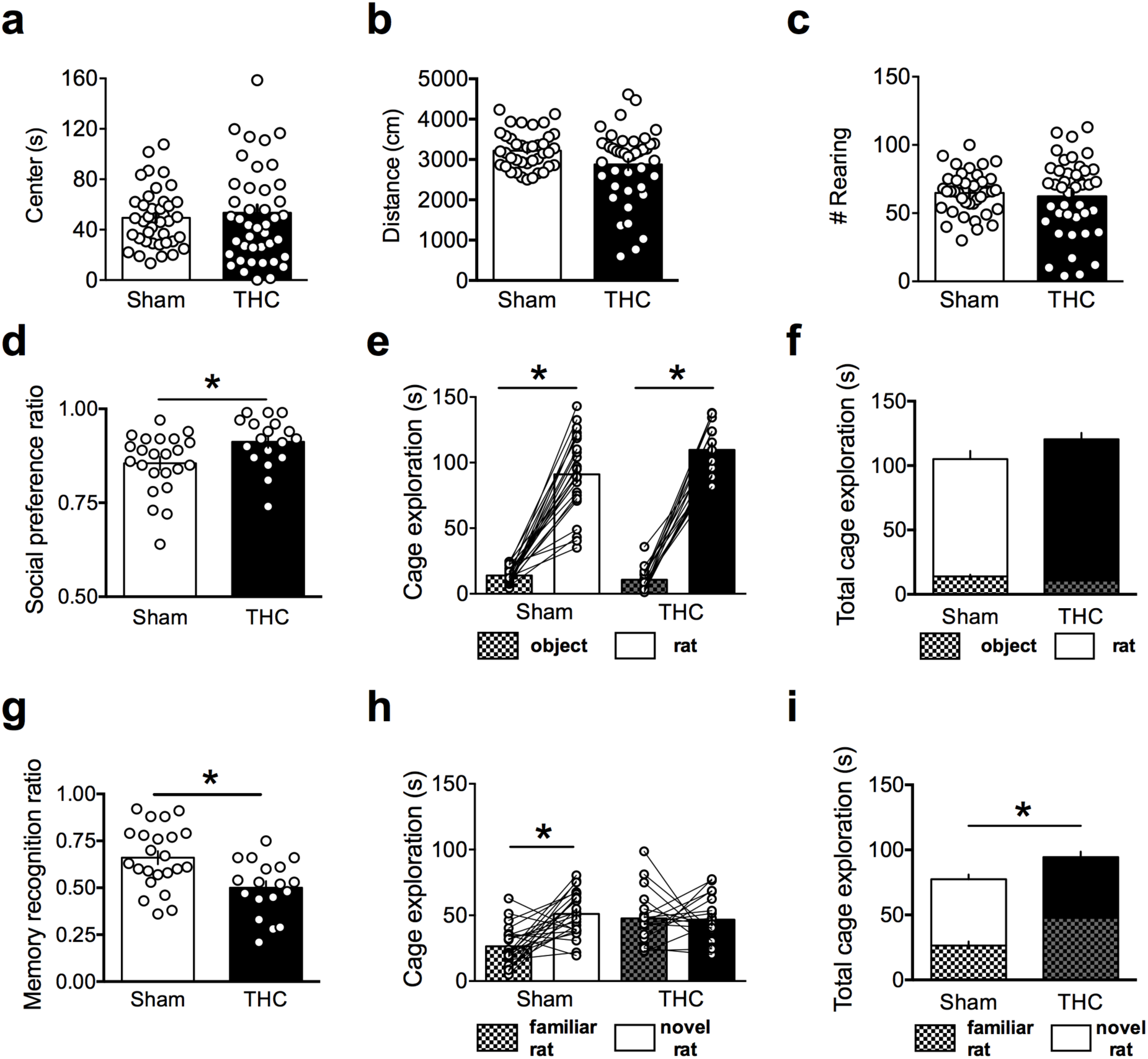
Perinatal THC exposure alters social approach and memory without changing behavior in the open field environment. **a-c**: Behaviors in the open field arena were not altered in the adult offspring of THC-treated as compared to those from sham-treated dams (N=42, 40 respectively). Specifically, no differences were found in the total distance covered during the trial, the bouts of rearing behavior, nor in the time spent in the center of the arena (Two-tailed T-test, P = 0.1846, 0.9644 and 0.7765, respectively). **d-f**: Social discrimination between an object and a novel rat is enhanced in the adult offspring of THC-treated, as compared to sham-treated, dams (N=18, 23 respectively). **d**: Specifically, the social preference ratio is significantly higher in the offspring of THC-treated dams (Two-tailed T-test, P = 0.0130). **e**: Both groups showed significant preference for the novel rat over the novel object (One-way ANOVA, F_3,78_ = 137.2; Tukey’s post-hoc analysis, P<0.0001 for both groups). **f**: The total time spent exploring the novel object and the novel rat did not differ between groups (Two-tailed t-test, P = 0.1518). **g-i**: In the subsequent social memory test, the adult offspring of THC-treated dams failed to discriminate between the familiar and novel rats. **g**: The social memory ratio is significantly higher in the offspring of sham-treated dams as compared to THC-treated dams (Two-tailed t-test, P = 0.0048). **h**: The offspring of sham-treated dams showed a significant preference for the novel partner rat (P<0.0001, Tukey’s post-hoc analysis following a significant One-way ANOVA, F_3,78_ = 9.016, P<0.0001). Conversely, no preference between the familiar and novel rats was demonstrated by the offspring of THC-treated dams (P = 0.9984, Tukey’s post-hoc analysis). **i**: The total time spent exploring non-discriminated rats was significantly higher in the THC group (Two-tailed t-test, P = 0.0086). *P<0.05

Next, based on previous findings ^26^ from our lab which have revealed sex-specific deficits in social interaction following *in utero* exposure to cannabinoids, we tested the adult offspring of dams given THC during the early postnatal period in a two-stage social task to assess both social preference/approach (Figure 1d-f) and social memory (Figure 1g-i).

Here, we found that during the initial testing period, the adult offspring of sham- or THC-treated dams both exhibited a significant preference for exploring the novel rat rather than the novel object, as determined by time spent at the two cages containing the social stimulus (Figure 1e). Interestingly, this preference, as determined by a ratio between the time spent at these two sites, unveiled a stronger social preference amongst THC-exposed rats, as compared to the sham group (Figure 1d). No significant difference was noted in the total time spent exploring both cages (Figure 1f). Therefore, both sham- and THC-exposed rats exhibit significant social preference over novel object stimuli, a behavioral characteristic which appears to be augmented in the latter group.

Next, in the social memory task, sham-exposed rats displayed the expected preference for a novel rat over the familiar rat from the first testing period (Figure 1g-h). However, THC-exposed rats failed to exhibit such a preference, as revealed by equivalent amounts of time spent exploring the cages containing both the novel and familiar stimulus rats (Figure 1h). Interestingly, the total time spent exploring non-discriminated rats was significantly higher in the THC-exposed rats, as compared to the sham group (Figure 1i). This significant difference owes to increased time spent on the familiar rat, without a lower amount of time spent on the novel rat, as compared to these times of exploration exhibited by the sham-exposed rats. Thus, we conclude that perinatal THC exposure increases overall social exploration at the cost of reducing discrimination between novel and familiar social stimuli. Sex-specific details can be found in Table 2.

### Perinatal THC exposure alters multiple forms of synaptic plasticity in the PFC at adulthood

The PFC plays a significant role in social behavior ^36^. Previously, we have shown that deficits in social behavior such as those described here are correlated with alterations to synaptic plasticity in the PFC ^26^ including a loss of eCB-mediated long-term depression (LTD) induced by a 10-minute, 10Hz stimulation at excitatory synapses onto layer 5 PFC pyramidal cells. Thus, we next investigated this form of LTD in the PFC of adult rats exposed to THC during the early perinatal period (Figure 2).

**Figure 2.**
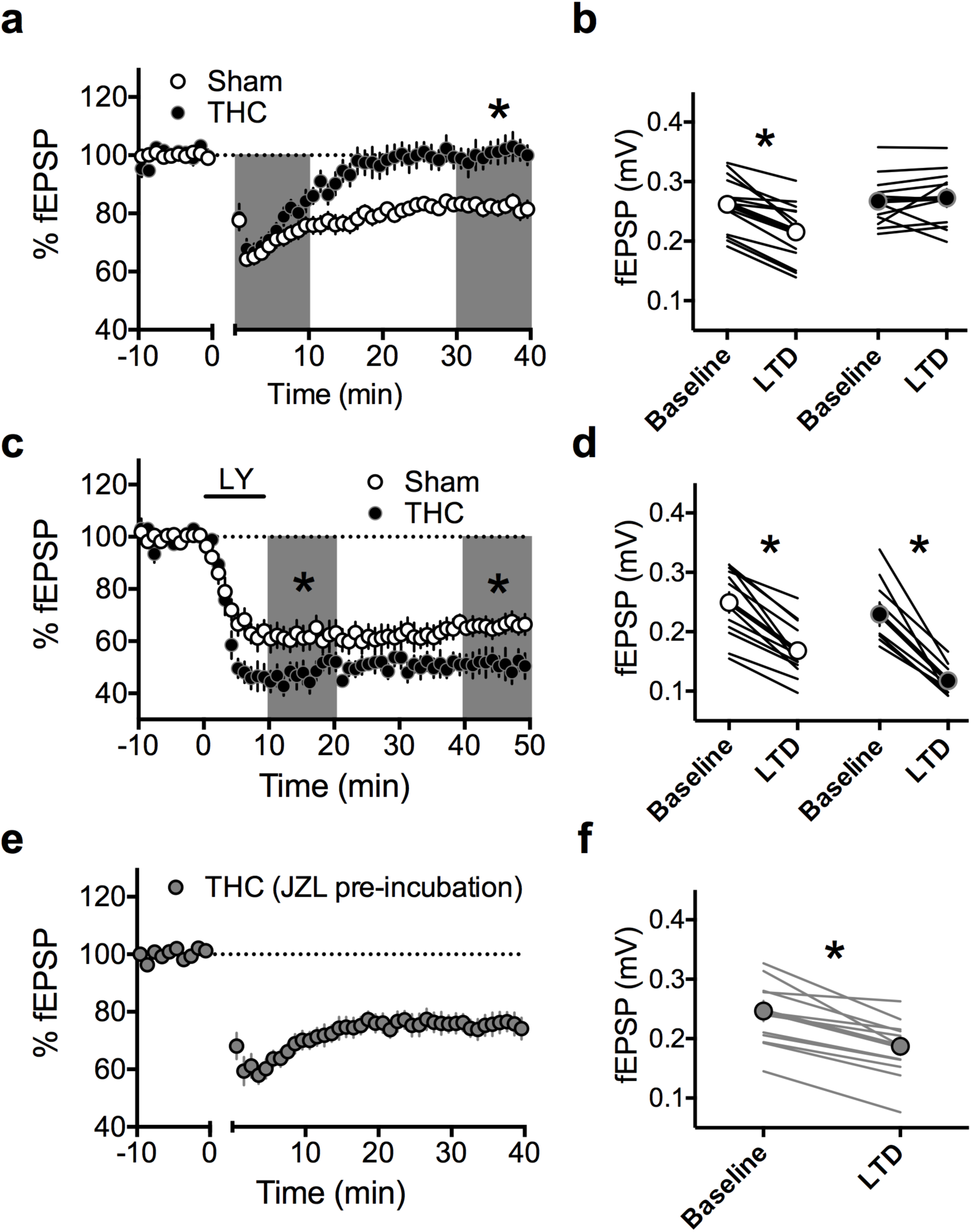
Perinatal THC exposure induces a selective deficit in LTD in the PFC of adult offspring. **a**: A 10-minute, 10Hz field stimulation of layer 2/3 cells in the PFC of the adult offspring of vehicle-treated dams (N=14) elicited a robust eCB-LTD at deep layer synapses. However, this same protocol failed to induce eCB-LTD in the adult offspring of dams treated with THC (N=11). No differences were found during the initial post-tetanus period (0-10 minutes post-tetanus), however at 30-40 minutes post-tetanus, the baseline-normalized %fEPSP was significantly lower in sham-exposed, as compared to THC-exposed rats (Two-way RM ANOVA, F_1,23_ = 10.63, P = 0.0034. Sidak’s multiple comparisons test, P = 0.3972 and P<0.0001 for 10 minutes and 40 minutes post-tetanus, respectively). **b**: fEPSP magnitude at baseline (−10 to 0 minutes) and LTD (30-40 minutes post-tetanus) values corresponding to the normalized values in **a** (Two-way RM ANOVA, F_1,23_ = 20.25, P = 0.0002. Sidak’s multiple comparisons test, P<0.0001 and P = 0.7795 for sham and THC, respectively). **c**: LTD mediated by mGlu_2/3_ receptors (mGluR-LTD) is augmented in THC-exposed offspring. mGluR-LTD, induced via a 10-minute application of LY379268 (LY; 30nM), produced a significant depression at deep layer synapses of the PFC in the adult offspring of both sham- and THC-treated dams (N=12 and 8, respectively). During both the first ten-minutes post-drug and the last ten-minutes (i.e. 30-40 minutes post-drug), fEPSP depression was significantly larger in THC-exposed offspring, as compared to sham-exposed (Two-way RM ANOVA, F_1,18_ = 7.913, P = 0.0120. Sidak’s multiple comparisons test, P = 0.0068 and P = 0.0048 at 10-minutes and 40-minutes, respectively). **d**: fEPSP magnitude at baseline (−10 to 0 minutes) and LTD (30-40 minutes post-drug) values corresponding to the normalized values in **c** (Two-way RM ANOVA, F_1,18_ = 105.5, P<0.0001. Sidak’s multiple comparisons test, P<0.0001 for both sham and THC). **e**,**f**: Enhancing 2-AG levels restores eCB-LTD in THC-exposed progeny. Following a >45-minute pre-incubation with the MAGL inhibitor, JZL 184 (4µM), the previously ineffective 10-minute, 10Hz protocol effectively induced a significant eCB-LTD at deep layer synapses of PFC slices obtained from the adult offspring of THC-treated dams (N=11; Two-tailed t-test, P = 0.0084). *P<0.05

First, we established that a 10-minute, 10Hz stimulation of superficial-layers of the PFC in slices obtained from the adult offspring of sham-injected dams induces a robust LTD at layer 5 synapses (Figure 2a-b). However, this same protocol failed to elicit LTD in slices obtained from THC-exposed animals. Thus, we can conclude that perinatal THC exposure induces a lasting deficit in eCB-LTD in the PFC.

To determine if this ablation of LTD in the PFC was limited to eCB-LTD or a more global impact, we next examined a distinct form of LTD in the PFC mediated by mGlu2/3 receptors ^37,38^ which has previously been shown to be disrupted by chronic exposure to drugs of abuse ^39^ but not by in utero cannabinoid exposure ^26^. Previous work has shown that mGlu2/3 LTD and eCB-LTD can mutually occlude each other in the nucleus accumbens 40. Further, shared pools of G_i/o_, normally sequestered by CB1R ^41^, are likely made more available as a result of CB1R desensitization/ablation of eCB-LTD, thereby permitting enhanced mGlu2/3-LTD ^40^. We therefore sought to test this possibility in our THC-exposed offspring by exposing PFC slices for 10-minutes to the mGlu2/3 agonist, LY379268 (300nM) in order to induce an mGlu2/3-dependent LTD. As predicted, this drug application elicited a significant LTD at layer 5 synapses in slices obtained from the adult offspring of sham-treated dams (Figure 2c-d). Similarly, the 10-minute application effectively elicited LTD in slices obtained from THC-exposed rats. Interestingly, the magnitude of this mGlu2/3 LTD is significantly larger in the THC-exposed rats, as compared to sham-exposed rats. Previously, a similar compensatory enhancement of mGlu2/3 LTD has been noted in mice chronically exposed to THC wherein eCB-LTD is abolished ^40^. Thus, perinatal THC exposure appears to abolish eCB-LTD while enhancing mGlu2/3-LTD in the PFC of adult rats.

eCB-LTD requires the participation of endocannabinoids, particularly 2-AG in the PFC ^32^. We sought to determine if enhanced levels of 2AG could benefit to eCB-LTD in THC exposed rats. Thus, PFC slices were incubated in JZL184 (4 µM, a potent inhibitor of Monoacyl-glycerol lipase the main enzyme degrading 2AG) in order to increase basal 2-AG levels prior to the 10-minute, 10Hz stimulation. Here, slices obtained from the adult offspring of THC-treated dams were found to exhibit robust, lasting LTD at layer 5 synapses following this pre-incubation. These data therefore indicate that increased levels of 2-AG in the PFC effectively restores eCB-LTD under these conditions.

### Perinatal THC abolishes TBS-LTP in the PFC at adulthood

Next, we elected to investigate additional forms of plasticity in the PFC of the adult offspring of THC-treated dams in order to determine the extent of altered PFC plasticity. Cannabinoids induced perturbations go beyond eCB-mediated synaptic plasticity ^42,43^. Notably, a single cannabinoid exposure abolishes theta-burst induced long-term potentiation (TBS-LTP) in the PFC of adult male rats ^30^. In order to investigate whether a similar perturbation occurred in the PFC of adult, THC-exposed offspring, a TBS was applied to superficial layers of acute PFC slices obtained from these rats during simultaneous recording of extracellular field EPSPs at deep layer synapses. Interestingly, while this TBS protocol effectively induced a lasting synaptic potentiation in slices obtained from the adult progeny of sham-treated dams, no such plasticity was observed in slices obtained from THC-treated dams (Figure 3a-b). As with the loss of eCB-LTD, we sought to determine if increasing basal levels of endocannabinoids would mechanistically ameliorate this loss of LTP. Thus, slices were incubated with JZL184, for >45 minutes prior to recording. Here, the TBS protocol effectively elicited a strong and lasting synaptic potentiation (Figure 3c-d). Thus, we observed that, as with the lost eCB-LTD, increased basal levels of 2-AG effectively restores lost TBS-LTP in the PFC of adult, THC-exposed rats.

**Figure 3.**
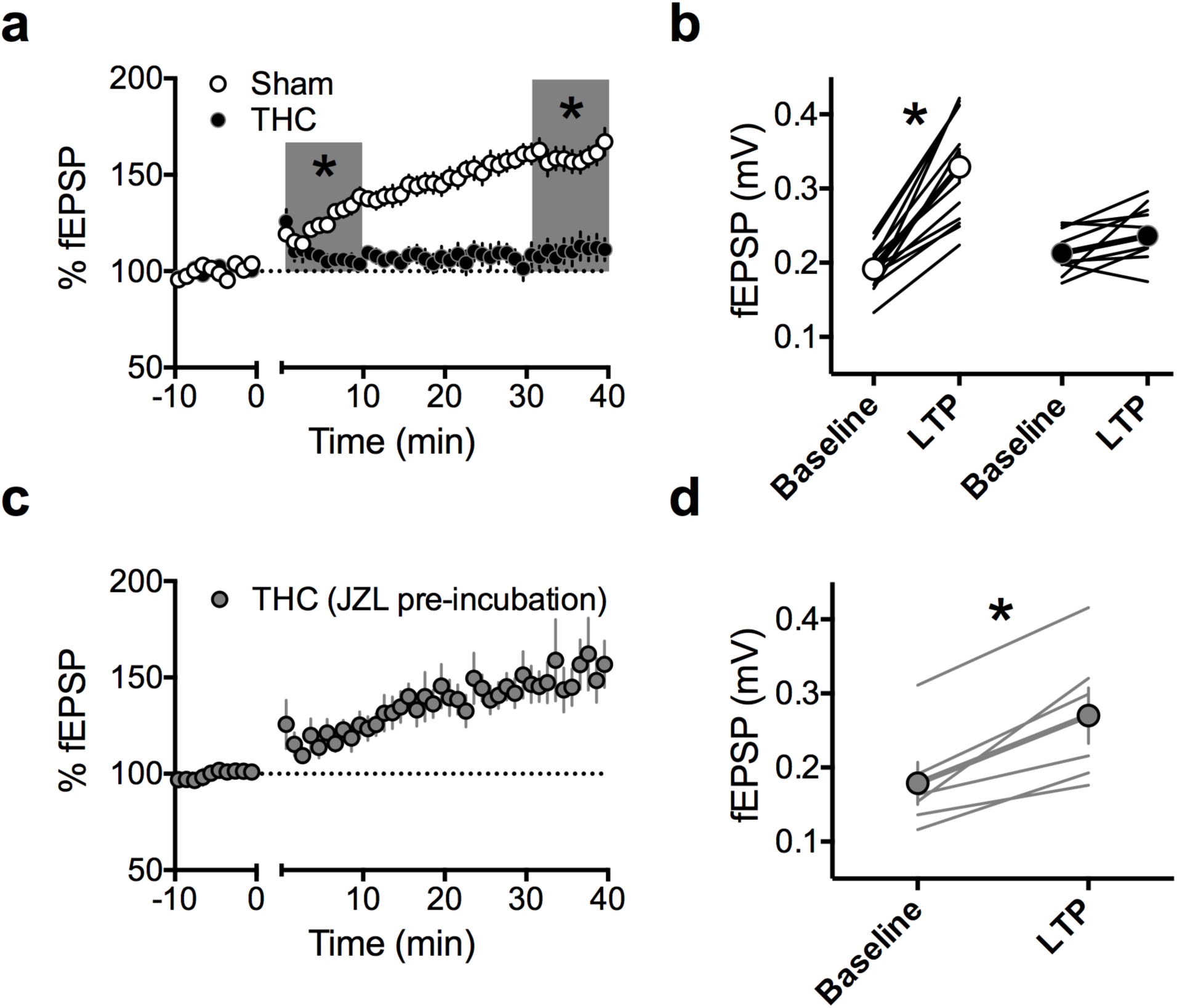
Perinatal THC exposure abolishes theta-burst stimulation (TBS) LTP in the PFC of adult offspring in a 2-AG-dependent manner. **a**: A TBS protocol (5 pulses at 100hz, repeated 4 times) at layer 2/3 cells in the PFC of the adult offspring of sham-treated dams (N=13) elicited a robust LTP at deep layer synapses. However, this same protocol failed to induce LTP in PFC slices obtained from the adult offspring of THC-treated dams (N=10). During both the first ten minutes post-tetanus and the last ten minutes of recording (i.e. 30-40 minutes post-tetanus), the normalized %fEPSP was significantly larger in slices obtained from sham-exposed rats, as compared to those of THC-exposed rats (Two-way RM ANOVA, F_1,21_ = 32.57, P<0.0001. Sidak’s multiple comparisons test, P = 0.0094 and P<0.0001 for 10 minutes and 40 minutes post-tetanus, respectively). **b**: fEPSP magnitude at baseline (−10 to 0 minutes) and LTP (30-40 minutes post-tetanus) values corresponding to the normalized values in **a** (Two-way RM ANOVA, F_1,21_ = 27.34, P<0.0001. Sidak’s multiple comparisons test, P<0.0001 and P = 0.3125 for sham and THC, respectively). **c**,**d**: Enhancing 2-AG levels permits the subsequent induction of TBS-LTP in THC-exposed progeny. Following a >45-minute pre-incubation with the MAGL inhibitor, JZL 184, the previously ineffective TBS protocol effectively induced a robust LTP at deep layer synapses of PFC slices obtained from the adult offspring of THC-treated dams (N=6, Two-tailed T-test, P = 0.0085). *P<0.05

### Perinatal THC alters select parameters of cell-excitability in the PFC at adulthood

In utero cannabinoid exposure has previously been shown to alter basic parameters of cell-excitability in the PFC at adulthood ^26^. Thus, we next performed patch-clamp recordings of deep-layer PFC pyramidal neurons in acute PFC slices obtained from the adult progeny of either sham- or THC-treated dams. An input-output curve revealed no significant differences between these groups (Figure 4a), however a closer examination of current-injection induced action potentials revealed two key differences. First, the number of spikes elicited by current injections between 0-600pA was significantly lower in THC-exposed offspring, as compared to sham controls (Figure 4b). This effect was expectedly correlated with a significantly higher rheobase in these cells (Figure 4c). However, no change in the resting membrane potential was noted (Figure 4d), indicating that rather than being less excitable due to lower resting membrane values, other mechanisms drive the decreased excitability observed in these animals.

**Figure 4.**
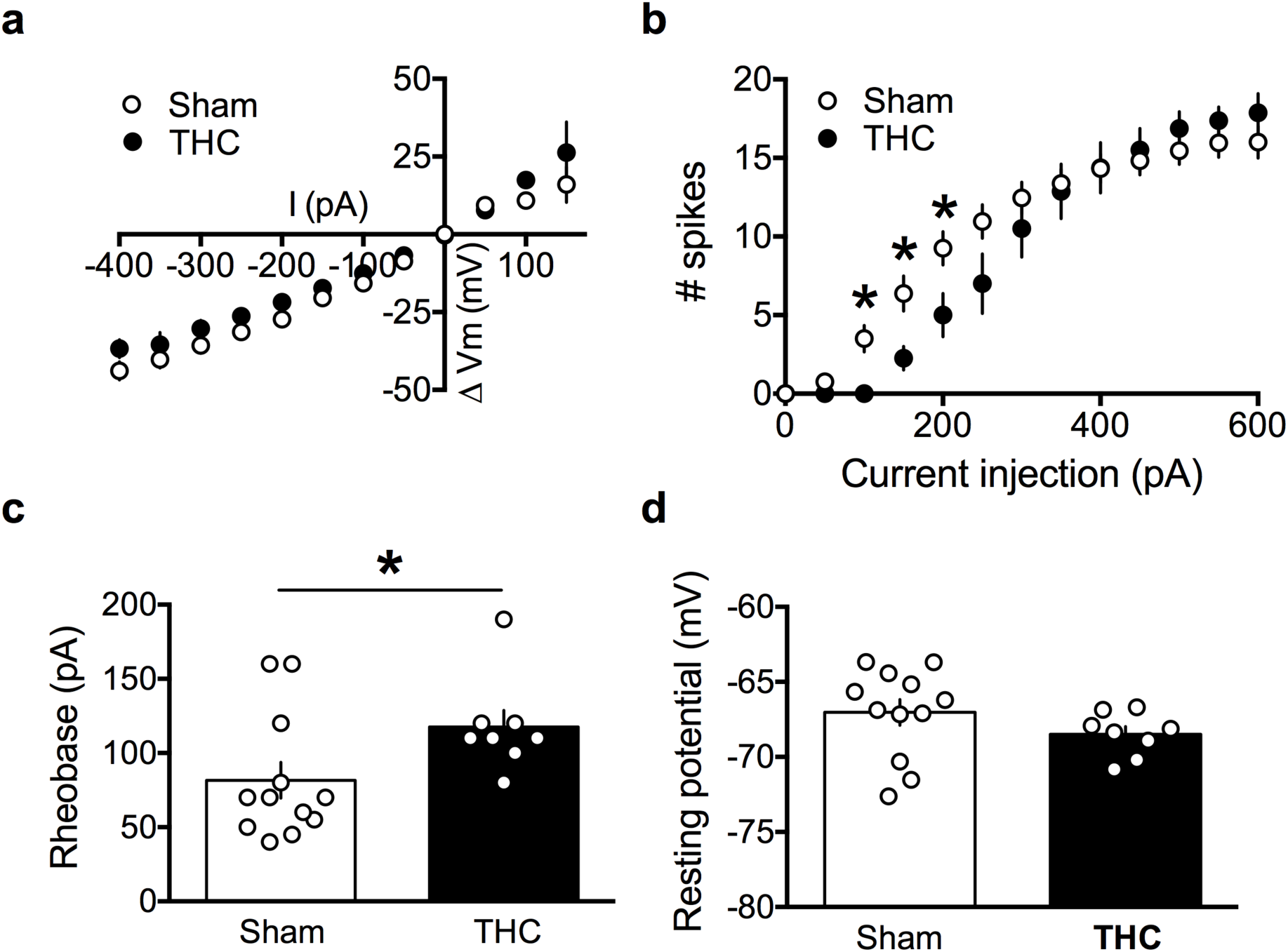
Perinatal THC exposure alters properties of intrinsic excitability of deep layer pyramidal neurons in the PFC of adult offspring. **a**: Current injection steps of 50pA from −400pA to 150pA revealed no differences in the I-V relationship in pyramidal neurons of the PFC between the adult offspring of sham- and THC-treated dams (N=12, 8 respectively). **b**: Action potentials elicited by progressive current injections from 0-600pA revealed a reduced number of spikes at low current injections in pyramidal neurons of the PFC in slices obtained from the adult offspring of THC-injected dams as compared to those from sham-treated dams. Significantly lower numbers of action potentials were observed following current injections of 100, 150 and 200pA (N=8, 12 respectively; Two-way RM ANOVA, F_20,360_ = 3.912, P<0.0001. Sidak’s multiple comparisons test, *P<0.05). **c**: Progressive current injections in 10pA steps from 0-200pA revealed that the minimum current injection required to elicit an action potential (i.e. rheobase) was significantly higher in deep layer pyramidal neurons of PFC slices obtained from the adult offspring of THC- as compared to sham-treated dams (N=8, 12 respectively; Two-tailed t-test, P = 0.047). d: No difference was observed in the resting membrane potential of deep layer pyramidal cells in PFC slices obtained from the adult offspring of THC-treated dams, as compared to those obtained from sham-treated dams (N=8, 12 respectively; Two-tailed t-test, P = 0.1646). *P<0.05

## Discussion

Here, we have discovered that perinatal exposure to THC via lactation induces significant behavioral and electrophysiological alterations lasting into adulthood. Specifically, we found that this THC exposure alters social behavior, as well as synaptic plasticity and basic parameters of cell-excitability in the PFC of adult male and female offspring of dams given THC during the first 10 days of postnatal life.

First, our behavioral explorations revealed that perinatal THC exposure augments social exploration at the cost of discrimination between novel and social interactions. Interestingly, these results diverge from those found following in utero THC exposure, wherein social exploration has been found to be reduced ^26,35^. Further, while in utero THC exposure appears to alter social behavior in a sexually divergent manner ^26^, here we have observed no differences between male and female progeny perinatally exposed to THC. The timing-specificity of long-term effects of drug exposure with regards to sexually dimorphic outcomes has been noted with several other substances, including tobacco ^44^, the anesthetic drug isoflurane ^45^ and toxins such as lead ^46^ and bisphenol-A ^47^.

At a systems level, we observed multiple alterations to synaptic plasticity in the PFC. First, eCB-mediated LTD was entirely absent at deep layer PFC synapses in slices obtained from the progeny of THC-treated dams. This finding is in line with previous findings from our lab demonstrating a loss of eCB-LTD in the PFC of in utero exposed offspring at adulthood ^26^ and extends upon previous findings that defects in PFC plasticity correlate with social dysfunction in other contexts, such as a mouse model of dietary polyunsaturated fat imbalance ^33^. Further, the loss of eCB-LTD correlated with an enhanced magnitude of mGlu2/3-LTD in the THC-exposed progeny. We surmise that this homeostatic compensation engages in order to maintain a functional working range of plasticity in the PFC and may result from increased availability of Gi/o messenger proteins which, prior to the loss of eCB-LTD, are likely to be sequestered by CB1 receptors ^41^ and therefore made less available for other forms of inhibitory plasticity, such as mGlu2/3-LTD.

The reappearance of eCB-LTD and TBS-LTP in the presence of enhanced 2-AG levels achieved via pre-incubation with the potent and selective MAGL inhibitor, JZL184, may owe to a previously demonstrated relationship between presynaptic NMDA receptors and CB1R which have been shown to detect co-incident pre- and post-synaptic activation during certain forms of both LTD and LTP at deep layer synapses of the PFC ^48^. Interestingly, elsewhere in the brain, NMDAR-LTD in the nucleus accumbens has also been found to require the participation of CB1R ^49^. Here, NMDAR-LTD is abolished following repeated cannabinoid self-administration and is later rescued through enhanced CB1R signaling achieved via the application of a CB1R positive allosteric modulator. Thus, the current data add to emerging evidence linking CB1R to NMDAR in the regulation of synaptic plasticity.

Finally, we observed that perinatal THC exposure decreases excitability of principle neurons of the PFC. As with other results of this study, this finding stands in contrast to those of earlier studies which have explored the consequences of in utero cannabinoid exposure, wherein increased excitability has been observed in the PFC ^26^. Rather, this finding mirrors those of chronic exposure to THC later in development, such as during the adolescent period wherein decreased excitability of PFC neurons has been observed in mice ^50^. Thus, we subscribe such differences to underlying state-dependent differences in the maturational window of development during which THC exposure takes place.

Together, our results indicate that perinatal THC exposure via lactation induces lasting deficits at multiple scales which persist into adulthood. THC-exposed offspring exhibit increased social exploration at the cost of discrimination, coupled with significant alterations to multiple forms of plasticity in the PFC which are normalized via enhanced basal 2-AG *ex vivo*. Increased excitability of principal neurons of the PFC may underlie or accompany these issues, and further investigations are required to further characterize the extent to which basic synaptic transmission may be impacted by this early life exposure. Further, both an increased breadth of behavioral investigations as well as extended characterizations of plasticity and synaptic functions in these animals in other brain regions are necessary to provide a more thorough picture of the extent to which perinatal cannabis exposure induces lasting deficits in brain function into adulthood.

## Supporting information

Scheyer 2020 Tables

## Author contributions

A.S., M.B. and O.J.M. designed research; A.S. and M.B. performed research; A.S. and M.B. analyzed data; A.S., M.B. and O.J.M wrote the paper. The authors declare no conflict of interest.

## Funding and Disclosures

This work was supported by the Institut National de la Santé et de la Recherche Médicale (INSERM); the INSERM-NIH exchange program (to A.F.S.); Fondation pour la Recherche Médicale (Equipe FRM 2015 to O.M.) and the NIH (R01DA043982 to O.M.).

## Declarations of interest

The authors declare no competing interests.

## Acknowledgements

The authors are grateful to the Chavis-Manzoni team members for helpful discussions.

