## Supplementary material for "Perinatal THC Exposure via Lactation Induces Lasting Alterations to Social Behavior and Prefrontal Cortex Function in Rats at Adulthood": Scheyer 2020 Tables

**Table 1. Open field data by sex**

| Condition | Test – Measure (unit) | Value | T-test (M v F) |
|---|---|---|---|
| Sham male | Open Field – Distance (cm) | 3120 ± 80.10<br>N=22 | P = 0.1490 |
| Sham female | Open Field – Distance (cm) | 3348 ± 130.8<br>N=17 | |
| THC male | Open Field – Distance (cm) | 2632 ± 190.9<br>N=28 | P = 0.0011 |
| THC female | Open Field – Distance (cm) | 3366 ± 78.81<br>N=14 | |
| Sham male | Open Field – Rearing (#) | 62.23 ± 3.365<br>N=22 | P = 0.2443 |
| Sham female | Open Field – Rearing (#) | 68.18 ± 3.730<br>N=17 | |
| THC male | Open Field – Rearing (#) | 57.71 ± 5,984<br>N=28 | P = 0.0598 |
| THC female | Open Field – Rearing (#) | 71.71 ± 4,050<br>N=14 | |
| Sham male | Open Field – Center (sec) | 54.85 ± 5.348<br>N=22 | P = 0.0839 |
| Sham female | Open Field – Center (sec) | 42.24 ± 4.674<br>N=17 | |
| THC male | Open Field – Center (sec) | 60.38 ± 8.744<br>N=28 | P = 0.0706 |
| THC female | Open Field – Center (sec) | 40.38 ± 6.293<br>N=15 | |

**Table 2. Social approach and social memory data by sex**

| Condition | Test – Measure (unit) | Value | T-test (M v F) |
|---|---|---|---|
| Sham male | Social Preference (ratio) | 0.8647 ± 0.01743 N=15 | P = 0.5089 |
| Sham female | Social Preference (ratio) | 0.8375 ± 0.03569 N=8 | |
| THC male | Social Preference (ratio) | 0.9375 ± 0.01268 N=12 | P = 0.0688 |
| THC female | Social Preference (ratio) | 0.8617 ± 0.03250 N=6 | |
| Sham male | Social Memory (ratio) | 0.6147 ± 0.03886 N=15 | P = 0.0876 |
| Sham female | Social Memory (ratio) | 0.7463 ± 0.05970 N=8 | |
| THC male | Social Memory (ratio) | 0.5433 ± 0.04061 N=12 | P = 0.0928 |
| THC female | Social Memory (ratio) | 0.4150 ± 0.05614 N=6 | |
| Sham male | Social Preference – Time exploring object (sec) | 14.20 ± 1.853 N=15 | P = 0.8121 |
| Sham female | Social Preference – Time exploring object (sec) | 13.61 ± 1.610 N=8 | |
| THC male | Social Preference – Time exploring object (sec) | 7.683 ± 1.587 N=12 | P = 0.1099 |
| THC female | Social Preference – Time exploring object (sec) | 16.77 ± 4.608 N=6 | |
| Sham male | Social Preference – Time exploring rat (sec) | 95.38 ± 8.016 N=15 | P = 0.3569 |
| Sham female | Social Preference – Time exploring rat (sec) | 83.02 ± 10.25 N=8 | |
| THC male | Social Preference – Time exploring rat (sec) | 116.4 ± 6.441 N=12 | P = 0.0182 |
| THC female | Social Preference – Time exploring rat (sec) | 96.40 ± 4.024 N=6 | |
| Sham male | Social Memory – Time exploring familiar rat (sec) | 29.31 ± 3.733 N=15 | P = 0.2200 |
| Sham female | Social Memory – Time exploring familiar rat (sec) | 20.90 ± 5.375 N=8 | |
| THC male | Social Memory – Time exploring familiar rat (sec) | 44.77 ± 5.473 N=12 | P = 0.4691 |
| THC female | Social Memory – Time exploring familiar rat (sec) | 53.59 ± 10.24 N=6 | |
| Sham male | Social Memory – Time exploring novel rat (sec) | 47.06 ± 4.528 N=15 | P = 0.1116 |
| Sham female | Social Memory – Time exploring novel rat (sec) | 58.58 ± 5.159 N=8 | |
| THC male | Social Memory – Time exploring novel rat (sec) | 52.09 ± 5,176 N=12 | P = 0.0638 |
| THC female | Social Memory – Time exploring novel rat (sec) | 36.04 ± 5.938 N=6 | |

**Table 3. LTD data by sex**

| Condition | Test – Measure (unit) | Value | T-test (M v F) |
|---|---|---|---|
| Sham male | eCB-LTD – Normalized fEPSP (30-40min post-tetanus) | 80.61 ± 3.408<br>N=8 | P = 0.5478 |
| Sham female | eCB-LTD – Normalized fEPSP (30-40min post-tetanus) | 83.90 ± 4.061<br>N=6 | |
| THC male | eCB-LTD – Normalized fEPSP (30-40min post-tetanus) | 104.4 ± 4.605<br>N=7 | P = 0.4821 |
| THC female | eCB-LTD – Normalized fEPSP (30-40min post-tetanus) | 97.26 ± 7.866<br>N=4 | |
| Sham male | mGlu2/3-LTD – Normalized fEPSP (30-40min post-drug) | 66.26 ± 4.196<br>N=6 | P = 0.8390 |
| Sham female | mGlu2/3-LTD – Normalized fEPSP (30-40min post-drug) | 67.58 ± 4.700<br>N=6 | |
| THC male | mGlu2/3-LTD – Normalized fEPSP (30-40min post-drug) | 56.31 ± 2.443<br>N=5 | P = 0.1384 |
| THC female | mGlu2/3-LTD – Normalized fEPSP (30-40min post-drug) | 42.30 ± 6.192<br>N=3 | |
| THC male (JZL) | eCB-LTD – Normalized fEPSP (30-40min post-tetanus) | 78.12 ± 7.002<br>N=5 | P = 0.6373 |
| THC female (JZL) | eCB-LTD – Normalized fEPSP (30-40min post-tetanus) | 74.17 ± 3.814<br>N=6 | |

**Table 4. LTP data by sex**

| Condition | Test – Measure (unit) | Value | T-test (M v F) |
| --- | --- | --- | --- |
| Sham male | TBS-LTP – Normalized fEPSP (30-40min post-tetanus) | 185.6 ± 26.32 N=6 | P = 0.3101 |
| Sham female | TBS-LTP – Normalized fEPSP (30-40min post-tetanus) | 155.4 ± 6.180 N=7 | |
| THC male | TBS-LTP – Normalized fEPSP (30-40min post-tetanus) | 114.5 ± 11.62 N=5 | P = 0.7022 |
| THC female | TBS-LTP – Normalized fEPSP (30-40min post-tetanus) | 109.0 ± 7.205 N=5 | |
| THC male (JZL) | TBS-LTP – Normalized fEPSP (30-40min post-tetanus) | 158.9 ± 18.37 N=3 | P = 0.5576 |
| THC female (JZL) | TBS-LTP – Normalized fEPSP (30-40min post-tetanus) | 145.0 ± 11.44 N=3 | |

**Table 5. Intrinsic properties data by sex**

| Condition | Test – Measure (unit) | Value | T-test (M v F) |
|---|---|---|---|
| Sham male | Resting membrane potential (mV) | -67.95 ± 1.408 N=6 | P = 0.3080 |
| Sham female | Resting membrane potential (mV) | -66.11 ± 0.9624 N=6 | |
| THC male | Resting membrane potential (mV) | -67.49 ± 0.7115 N=3 | P = 0.1503 |
| THC female | Resting membrane potential (mV) | -69,09 ± 0.5940 N=5 | |
| Sham male | Rheobase (pA) | 86,67 ± 18.06 N=6 | P = 0.7003 |
| Sham female | Rheobase (pA) | 76.67 ± 17.64 N=6 | |
| THC male | Rheobase (pA) | 136.7 ± 26.67 N=3 | P = 0.3697 |
| THC female | Rheobase (pA) | 106.0 ± 7.483 N=5 | |
